# TypeAssembly: Copy number estimation and allele typing for haplotype assemblies

**DOI:** 10.1101/2025.09.13.675996

**Authors:** Hong-Sheng Lai, Yu-Shi Liu, Hsien-Chun Chiu, Ting-Yu Chang, Hsin-Fu Lin, Yi-Hsuan Tseng, Tsung-Kai Hung, Sheng-Kai Lai, Jacob Shujui Hsu, Chia-Lang Hsu, Ya-Chien Yang, Pei-Lung Chen, Chien-Yu Chen

## Abstract

Accurately annotating complex genes in the human genome, particularly from haplotype assemblies, remains a significant challenge. To overcome this, we developed TypeAssembly, a local alignment-based framework for copy number estimation and allele typing. Operating in two modes, mode-FASTA and mode-VCF, TypeAssembly can define alleles by either sequence or variant information. We successfully applied it to annotate 41 genes in the MHC locus, 17 *KIR* genes, and, for the first time, 15 pharmacogenes across 466 haplotype assemblies. This study establishes TypeAssembly as a robust method for accurately annotating complex genomic regions and provides an evaluation of existing gene annotations and callers.

## 1 Background

With the rapid advancement of high-throughput sequencing technologies, obtaining high-quality whole genome sequencing (WGS) data from short-read sequences has become increasingly accessible, enabling more comprehensive analysis of genetic variations. However, for certain genes with complex structures, such as those involving copy number variations and structural variations, allele typing and copy number estimation remain challenging. Moreover, there is no reliable benchmark dataset available to assess the performance of the currently developed tools. Recently, phased haplotype assembly data validated through short-read, long-read, and linked-read sequencing, such as datasets from the Human Pangenome Reference Consortium (HPRC) [1] and the Chinese Pangenome Consortium (CPC) [2], have become available. These high-quality haplotype assemblies provide researchers with valuable resources to better understand genes with complex structures. Accurate annotations of these assemblies can significantly expedite the development of tools for current short-read sequencing technologies.

Pharmacogenes are among the most structurally complex genes, with detailed clinical annotations provided by the Clinical Pharmacogenetics Implementation Consortium (CPIC) [3]. For instance, typing the *CYP2D6* gene presents significant challenges due to its copy number variations and hybrid gene structures involving its pseudogene, *CYP2D7*. Accurately identifying allele types in pharmacogenes is critical, as different allele types directly influence drug metabolism in the human body. Existing tools, such as PharmCAT [4], Aldy [5], and DRAGEN [6], are optimized for short-read WGS or targeted capture sequencing. However, the development of these tools faces limitations in lacking accurate information on copy numbers or complex structural variants, complicating the process of continually enhancing the accuracy of detection tools. While GeT-RM [7] has been a standard solution validated across multiple laboratories, its utility has diminished due to the rapid and continuous updates in pharmacogenomic databases such as PharmVar [8] and PharmGKB [9].

The major histocompatibility complex (MHC) gene locus and killer cell immunoglobulin-like receptor (KIR) genes play critical roles in immune function and are associated with a range of diseases, including autoimmune disorders, cancer, and infectious diseases. Their high polymorphism and structural complexity pose significant challenges for accurate annotation, prompting extensive research efforts focused on their characterization in human genome assemblies. Recently, Immuannot [10], a tool designed for full gene resolution of both human leukocyte antigen (HLA) and *KIR* genes, has been developed, demonstrating near-consensus results with SKIRT [11] in the annotation of 17 *KIR* genes. Additionally, a novel framework for *HLA* genotyping has been introduced [12], offering a comprehensive comparison with short-read *HLA* callers and successfully annotating 11 *HLA* genes.

In this study, we introduced TypeAssembly, a local alignment-based approach that performs allele typing on haplotype assemblies in two modes: mode-FASTA and mode-VCF. Using mode-FASTA, we annotated 94 haplotype assemblies from the HPRC first release, validating our results against consensus annotations for 17 *KIR* genes from the IPD-KIR database [13] and 41 genes in the MHC gene locus from the IPD-IMGT/HLA database [14] against Immuannot [10]. Through mode-VCF, we annotated 15 pharmacogenes listed in PharmVar, which previously lacked annotations. To demonstrate TypeAssembly’s utility in assessing short-read sequencing tools, we compared its results with those of DRAGEN and Aldy on 44 short-read WGS datasets. Above this, we annotate all above 73 complex genes on the latest HPRC second release, and extend the annotation to the extra 464 haplotype assemblies. By establishing the first comprehensive pharmacogene annotations, TypeAssembly paves the way for advancements in pharmacogenomics research. Furthermore, it provides a robust framework for annotating haplotype assembly data across a wide range of genes with well-defined allele information.

## 2 Results

### 2.1 Overview of TypeAssembly

The primary objective of TypeAssembly is to identify the locations of the gene regions within haplotype assemblies and analyze their genetic variations. There are two modes for running TypeAssembly owing to the different nature of allele definitions. For alleles defined in FASTA format, which often vary in length, such as in MHC and *KIR* regions, mode-FASTA is used (Fig. 1 (a)). TypeAssembly use wildtype sequences to identify the corresponding regions to get the target sequence of target genes. Then we align the target sequence against both the genomic sequence of alleles and the coding sequence of alleles, analyzing where the variants happen and determining the allele types or defining the novel alleles.

**Fig. 1:**
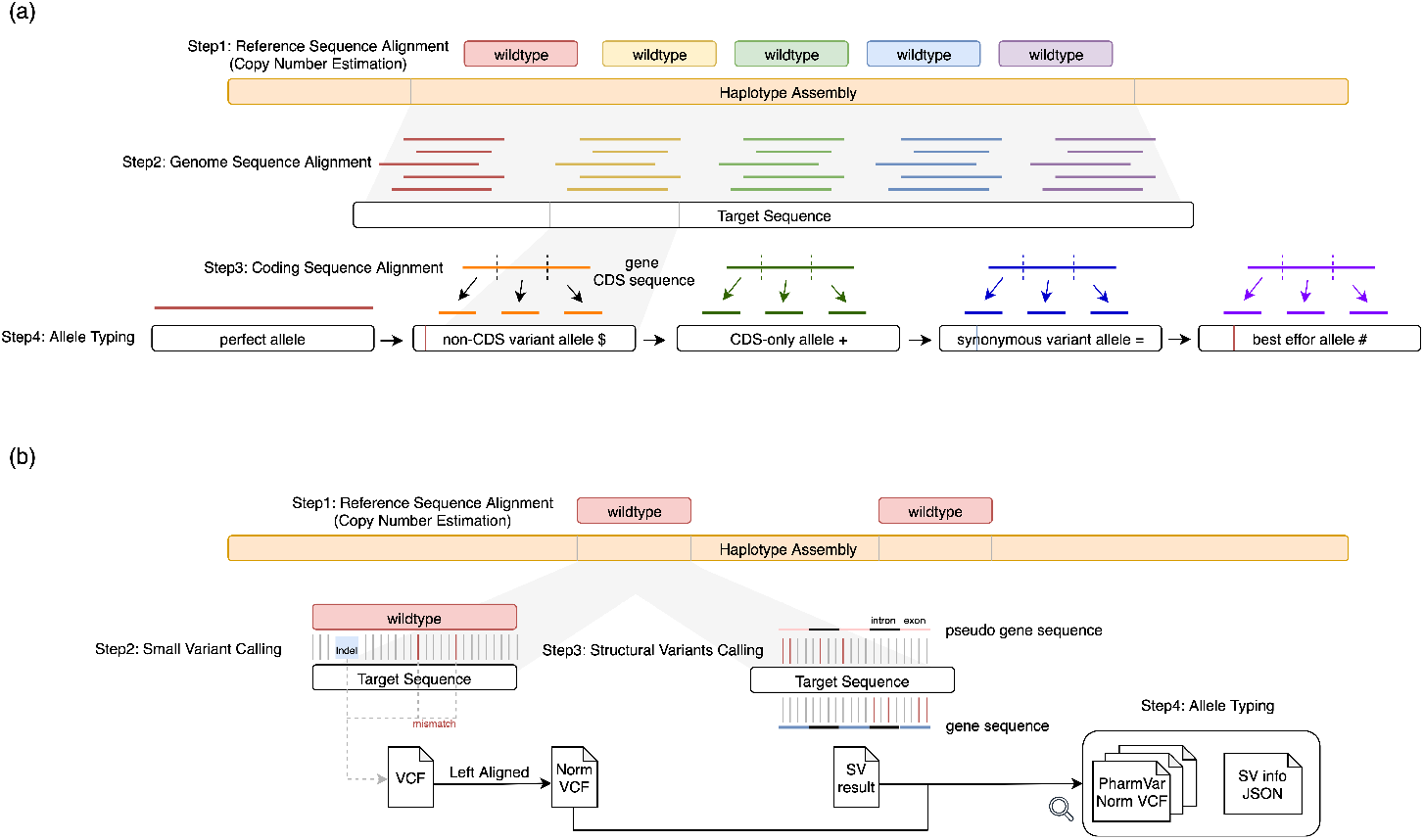
The overview of TypeAssembly pipelines. TypeAssembly has two running modes: mode-FASTA and mode-VCF. (a) In mode-FASTA, we align wildtype sequences to find the target sequence in the haplotype assembly. We then align a library of known genomic sequences of alleles and coding sequences of alleles against this target sequence. By analyzing where variants land, it can assign a known allele (perfect allele) or help characterize a novel allele (with $, +, =, # suffix). The allele typing should follow the order. (b) In mode-VCF, we also align the wildtype sequences to find the target sequence in the haplotype assembly. We then call small variants (e.g., indels and SNPs), and structural variants (SVs). Finally, it integrates the results to determine the final allele composition for each gene copy based on the PharmVar and the SV information.

If the alleles of a gene are defined by Variant Calling Format (VCF) format and have many non-deterministic variants, such as pharmacogenes, we may use mode-VCF (Fig. 1 (b)). First, TypeAssembly uses wildtype sequences to estimate the copy number of a target gene or to detect full gene deletions. The step is followed by extracting the corresponding gene sequences from the haplotype assembly data. When multiple copies of a target gene are identified, each copy is processed independently in subsequent steps. Next, we use the alignment from wildtype sequences and target sequences to identify small variants, such as single-nucleotide polymorphisms (SNPs) and small insertions or deletions (indels). The results are stored in VCF files. For genes with structural variants, TypeAssembly further analyzes the coding regions, incorporating exon and intron sequences from the gene and its pseudogene. This step allows us to identify regions where gene conversions or partial deletions have occurred. Finally, we perform allele typing using allele definitions provided in the VCF files and integrate structural variant results to determine the final allele composition.

### 2.2 Annotations for KIR and MHC

#### 2.2.1 Benchmark for HPRC First Release against Immuannot

Using wildtype sequences to estimate the region may have some risk for *KIR* and MHC, because the lengths are varied. The IPD-KIR database contained 1,548 genomic sequence alleles across 17 *KIR* genes. Among 1,548 alleles, we observed two alleles with alignment coverage below 50% relative to their corresponding wildtype sequence. To ensure TypeAssembly covered all possible regions, these two low-coverage alleles were incorporated into our custom *KIR* wildtype sequence sets, resulting in 19 wildtype sequences. Similarly, for the MHC gene locus, we started with the IPD-IMGT/HLA database, which comprised 24,897 genomic alleles across 41 genes. We identified and added 13 alleles with low reference coverage to our MHC wildtype sequence set, resulting in 54 wildtype sequences.

We validated TypeAssembly using the HPRC first release, which includes 94 haplotype assemblies from 47 individuals. In the *KIR* genes, our allele calls were compared against 892 allele annotations from Immuannot. TypeAssembly identified 890 alleles, achieving a 97% concordance rate with 866 matching calls. Of the 866 concordant calls, 122 represented allele name updates reflecting updates in the IPD-KIR database. The minor discrepancies were investigated and attributed to our different calling criteria. Specifically, two alleles were not called by TypeAssembly due to our filters requiring *>*97% sequence identity and *>*90% length coverage. The 24 mismatched calls primarily arose from our strict requirement for full-length allele identification, except for best effort allele, in contrast to Immuannot, which allowed assigning genotypes based on partial genomic sequence alignments.

In the MHC gene locus, we compared against 3452 allele annotations from Immuannot. TypeAssembly identified 3439 alleles, achieving a 98.13% concordance rate with 3375 matching calls, including 538 database updates. 13 not-called alleles are annotated as DRB1*new, which are filtered out in TypeAssembly. Other mismatches are mostly because of our length filter.

Further analysis of mismatches suggests that TypeAssembly can resolve previous annotation inaccuracies. For instance, in the HG00733 maternal haplotype, Immuan-not reported a novel allele, 3DL3*00500new. TypeAssembly revealed this to be a partial alignment at the genomic sequence level, and we could not identify full coverage of the corresponding CDS within the transcript, indicating the original call may have been based on alignments outside the target transcript region. In this sample, TypeAssembly demonstrates that ensuring the alignment from the genomic sequence level to the coding sequence level can help us make sure the specific alignment regions and enhance the allele typing accuracy. In HG03492 maternal haplotype, Immuan-not annotated DQA1*05:01:01:03. However, in TypeAssembly, we found out that the haplotype assembly has one more adenine in a consecutive repetitive adenine region.

#### 2.2.2 Annotation of All HPRC Samples

In the HPRC second release, 46 individuals (92 haplotype assemblies) are also included in the HPRC first release. Therefore, our final annotation was conducted on an expanded cohort of 233 individuals (466 haplotype assemblies), combining data from the HPRC first and second releases. This cohort includes 92 haplotype assemblies common to both releases, where HG03492 is only in the first release.

In *KIR* genes, a consistency check on these 92 overlapping haplotype assemblies revealed only four minor allele call differences between the data releases. Three of these differences involved a change between a perfect-match allele and a non-CDS variant of the same allele, while one involved a change between a best effort allele and a non-CDS variant allele. In the MHC gene locus, 33 differences were revealed, where 10 involved changes between a best effort allele and a non-CDS variant allele, and others are changes between a perfect-match allele and a non-CDS variant. All the above differences happened in the last field of their allele definitions.

Within the final 466-haplotype dataset, we identified 4,478 *KIR* alleles, of which 2,609 (58.3%) were novel. Also, 16460 MHC alleles were identified, and 5355 of them are novel. A detailed breakdown of the allele distribution for each gene can be found in Fig. 3. TypeAssembly also revealed 41 unique *KIR* gene haplotype structures. The ten most frequent structures accounted for 420 (90.1%) of all haplotype assemblies and are visualized in Fig. 2.

**Fig. 2:**
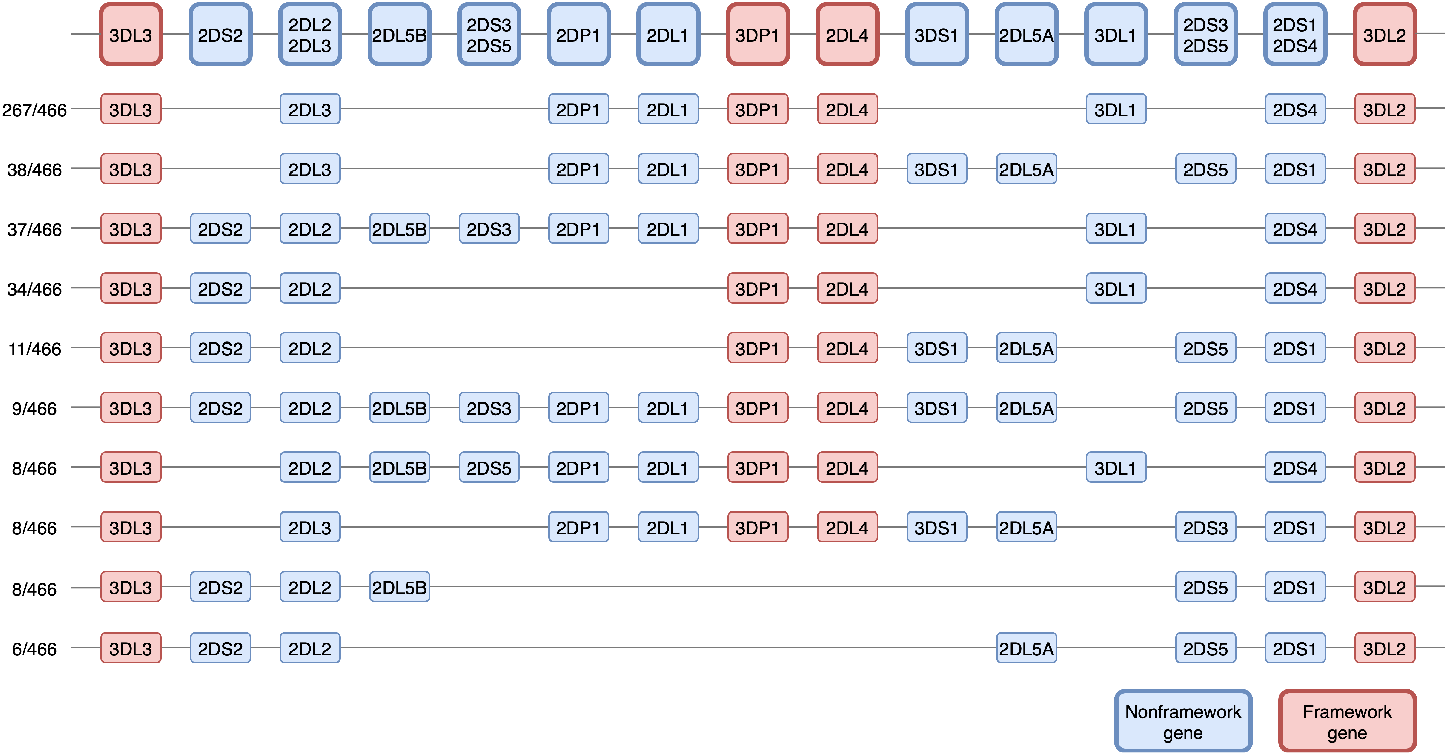
The top 10 common *KIR* gene haplotype structures.

**Fig. 3:**
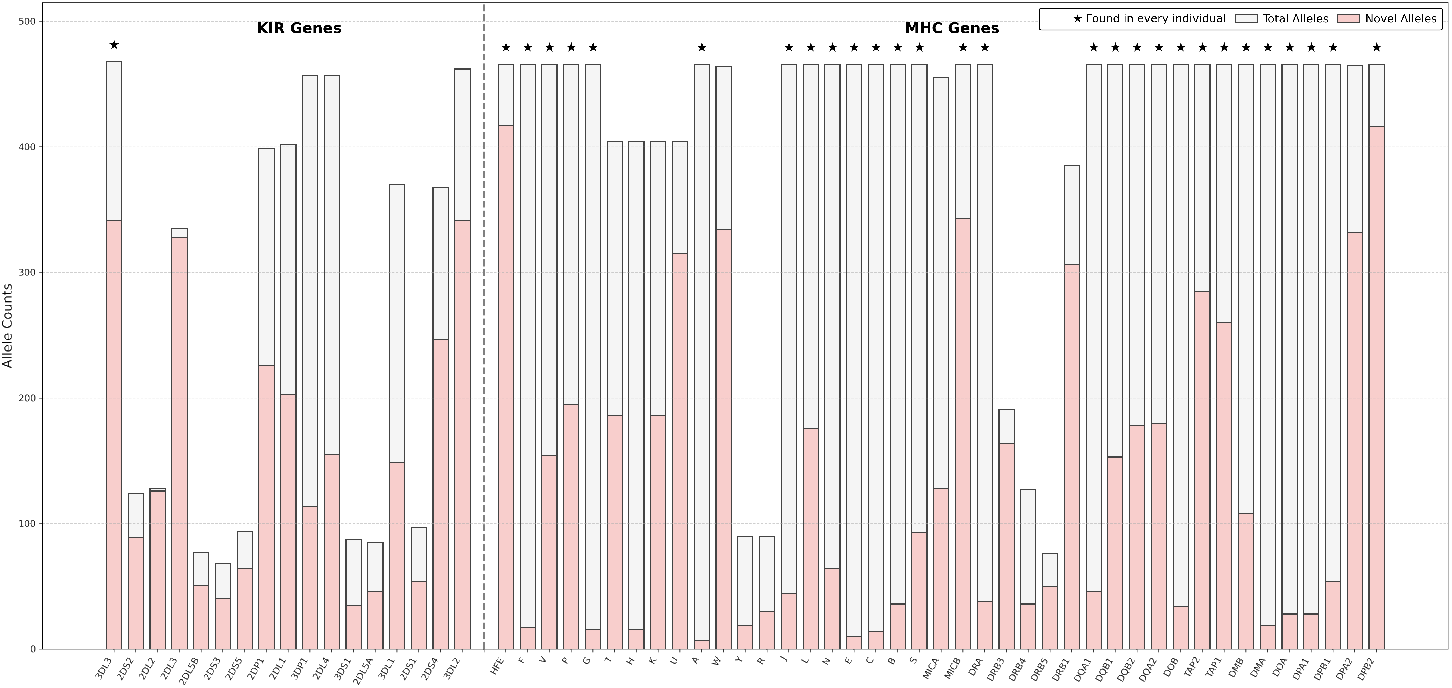
Distribution of (a) *KIR* and (b) MHC alleles and novel alleles.

In summary, TypeAssembly mode-FASTA provided an up-to-date annotation of *KIR* and MHC alleles for 466 HPRC haplotype assemblies. We demonstrated that TypeAssembly can not only be capable of handle complex gene regions with a properly provided FASTA file-based allele database, but also reduce errors that can arise from using partial alignment in the nearby genes to type the result.

### 2.3 Annotations for pharmacogenes

#### 2.3.1 Introduction to Current Tools

The objective of TypeAssembly mode-VCF is to accurately annotate the alleles on pharmacogenes and help current tools improve. Since there are no current annotations for these haplotype assembly data on pharmacogenes available, we directly compared 88 haplotype assemblies’ TypeAssembly annotation with the results from pharmacogenomics annotation tools using the WGS short reads released by HPRC first release. The evaluated short-read pharmacogenomics annotation tools include the commercially available DRAGEN pharmacogenomics caller (v4.2.4) and the non-commercial tool Aldy4.

#### 2.3.2 Mismatch

For the DRAGEN analysis, we focused on eight overlapping genes (Table 2) shared between the DRAGEN database and PharmVar. Certain overlapping genes, such as *NAT2*, were excluded due to substantial differences in the wildtype sequence used by DRAGEN. Additionally, *DPYD* was excluded from the comparison because it was called solely based on variants. This comparative result revealed potential errors not only in the DRAGEN pharmacogenomics caller for WGS data but also in the haplotype assemblies from HPRC (Fig. 4).

**Fig. 4:**
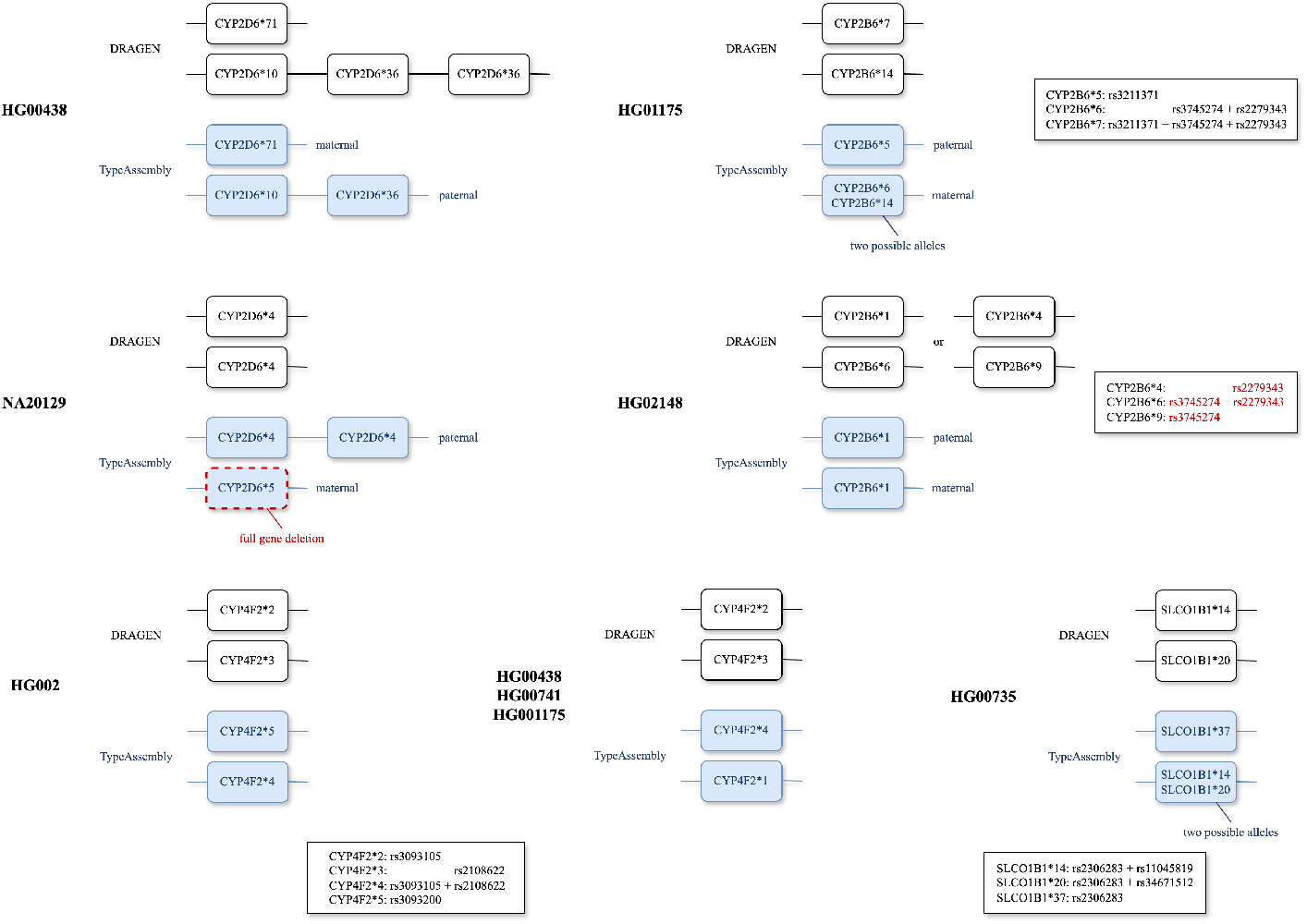
Mismatches between TypeAssembly and DRAGEN for *CYP2D6, CYP2B6, CYP4F2*, and *SLCO1B1* on 16 haplotype assemblies over 9 individuals. The red RSID indicates the missing variants in haplotype assemblies. The white box contains allele definitions to help the interpretation of phasing issues.

For example, in the HG00438 maternal strand, DRAGEN called *\*10+*36×2* for *CYP2D6*, whereas TypeAssembly identified only *\*10+*36* in the haplotype assembly data. Furthermore, using HiFi long-read data released by the HPRC first release, we cannot find the reads with two *\*36* copies. If the haplotype assembly is correct, this indicates a miscall by DRAGEN owing to missing phasing information. Similarly, in the NA20129 individual, DRAGEN reported *\*4/*4* for *CYP2D6*, but TypeAssembly revealed two copies of *\*4* (*\*4×2*) on the paternal strand and a full gene deletion on the maternal strand in the haplotype assemblies, indicating another miscall by DRAGEN. Using HiFi long-read data, we identified a read containing two copies of *CYP2D6*4* from NA20129, further supporting our findings. These two miscalls underscore the challenges associated with short-read data, particularly the difficulty in accurately estimating copy numbers based on read depth and the lack of phasing information compared to haplotype assembly data.

In the HG01175 individual, DRAGEN called *CYP2B6*7* (rs3211371 + rs3745274 + rs2279343) for the paternal strand. However, TypeAssembly identified rs3211371 on the paternal strand, while rs3745274 and rs2279343 were located on the maternal strand. This discrepancy alters the allele assignments: the maternal strand transitions from *CYP2B6*14* to two potential alleles, *CYP2B6*6* and *CYP2B6*14*, while the paternal strand is resolved as *CYP2B6*5*. We interpret this as a miscall by DRAGEN, again likely caused by the absence of phasing information.

In the HG02148 individual, DRAGEN called rs3745274 and rs2279343 in the CYP2B6 region, resulting in a determination of *\*1/*6* or *\*4/*9*. However, TypeAssembly did not identify these key variants in the haplotype assembly data. Notably, an analysis of HiFi long-read data reveals that 41% of alternative reads at the rs3745274 and rs2279343 positions are present, both located on the same strand. Based on this evidence, we conclude that the genotype is *\*1/*6*. This discrepancy suggests a potential assembly error. These findings highlight TypeAssembly’s ability not only to accurately call alleles in complex gene regions but also to detect possible errors in haplotype assemblies.

In the HG00735 individual, DRAGEN called *SLCO1B1*14* (rs2306283 + rs11045819) and *SLCO1B1*20* (rs2306283 + rs34671512). However, TypeAssembly identified that both rs11045819 and rs34671512 were located on the maternal strand. We interpret this as a miscall by DRAGEN, likely caused by the absence of phasing information.

In the *CYP4F2* gene, four individuals (HG002, HG00438, HG00741, and HG01175) exhibit phasing issues. DRAGEN assigns rs3093105 and rs2108622 to separate strands.

However, TypeAssembly identifies both variants as being on the same strand, changing the haplotype annotation from *\*2/*3* to *\*4/*N*. Here, N corresponds to *\*1* for HG00438, HG00741, and HG01175, while for HG002, N is identified as *\*5*, which DRAGEN is unable to call due to database differences.

For Aldy, we selected the same eight genes for comparison. The number of mismatches observed was higher than that with DRAGEN (Table 2), suggesting that our results align more closely with DRAGEN.

#### 2.3.3 Database Difference

Most pharmacogenomics tools rely on their own databases to store variant information. However, due to the continuous updates from PharmVar, maintaining standardization among these tools is challenging (Tables 2, 2). For example, in the individual HG02572, DRAGEN identifies the allele as *CYP2D6*1/*1*, whereas TypeAssembly identifies it as *CYP2D6*1/*152*. This discrepancy occurs because DRAGEN version 4.2.4 lacks a definition for the *\*152* allele. Similarly, for CYP4F2, DRAGEN v4.2.4 only recognizes alleles *\*1, *2*, and *\*3*, while PharmVar v6.1.6 now includes alleles *\*1* through *\*17*, resulting in nine discrepancies. These differences were also observed in Aldy.

In contrast, TypeAssembly simplifies this process by relying solely on VCF normalization from PharmVar rather than creating a dedicated database for small variant calling. This approach enables seamless integration of PharmVar updates without requiring extensive reconfiguration, making TypeAssembly a convenient tool for annotating alleles in pharmacogenes and other genes with well-defined variant information.

#### 2.3.4 Structural Variant Calling

To demonstrate the structural variant calling capability of TypeAssembly, we analyzed two hybrid structures from HPRC samples: *CYP2D6*36* from the HG005 paternal strand and *CYP2D6*68* from the HG00733 paternal strand. In *CYP2D6*36* (Fig. 5 (a)), a gene conversion occurs from *CYP2D6* to *CYP2D7* after exon 9. By aligning each exon and intron individually to the target sequence, we confirmed that the region surrounding exon 9 is more similar to *CYP2D7* (778/781 matches, 99% identity) than to *CYP2D6* (243/255 matches, 95% identity). The bar plots reveal significant differences in introns 2 and 6, which align more closely with *CYP2D6* in these regions. From the alignment results for *CYP2D7*, we observed that the first six base pairs (bp) of intron 3 and the entirety of intron 6 failed to align with the target sequences. Additionally, the first two bp of *CYP2D7* exon 1 and the last 526 bp of *CYP2D6* exon 9 align exclusively with the alternate gene, further supporting the hybrid nature of this structure. These findings confirm that *CYP2D6*36* is a hybrid structure that begins with *CYP2D6* and transitions to *CYP2D7*, with the conversion initiating at exon 9. For *CYP2D6*68* (Fig. 5 (b)), the gene conversion begins at the end of intron 1.

**Fig. 5:**
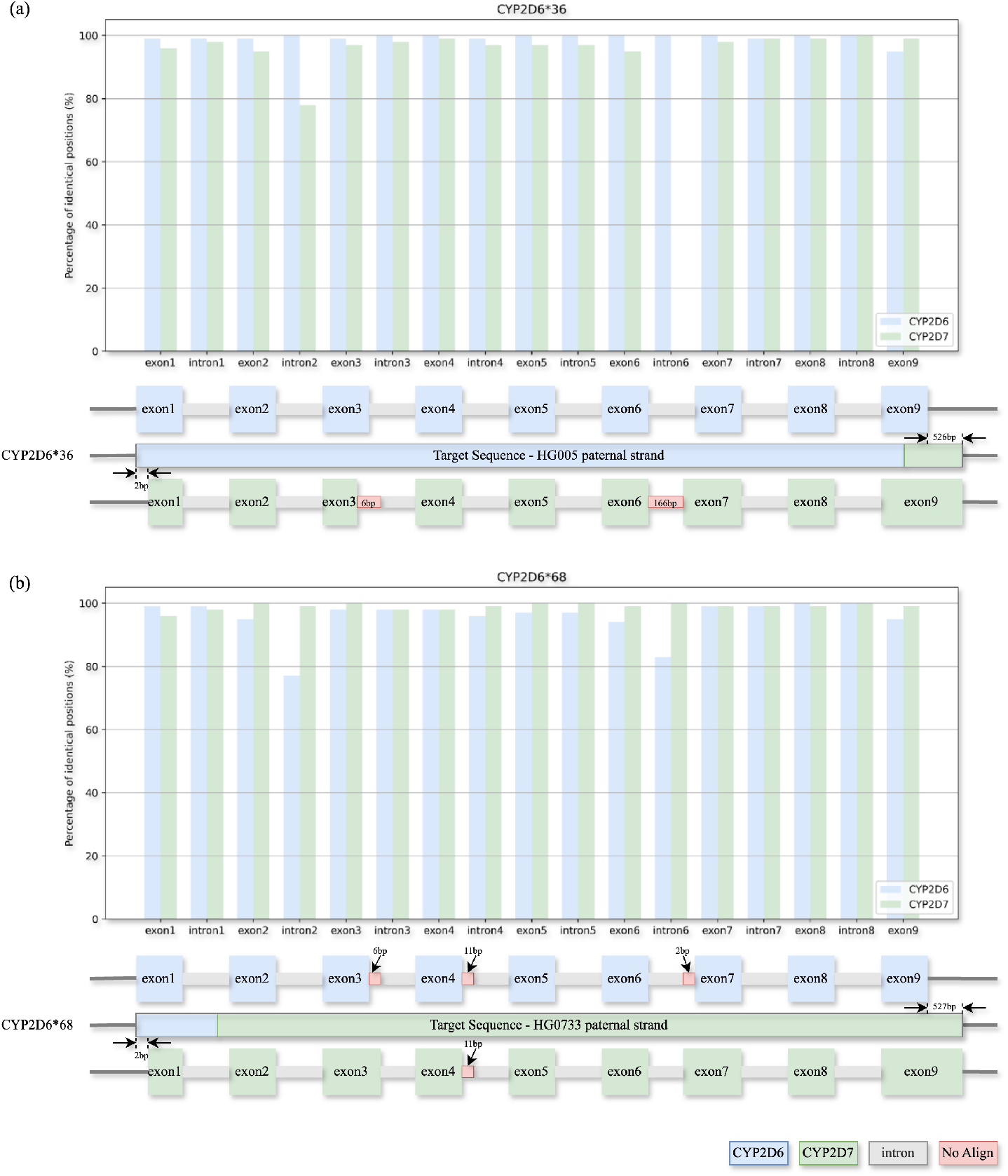
Gene conversions in *CYP2D6* observed on the target regions of HG005 and HG00733. Percentage of identical positions in exon and intron alignments between the *CYP2D6* /*CYP2D7* reference exon/intron sequences and the target sequence of (a) *CYP2D6*36* and (b) *CYP2D6*68*. Conversion for *CYP2D6*36* starts at exon 9, while the conversion of *CYP2D6*68* starts at intron 1.

Alignment analysis revealed that the target sequence after exon 2 is notably more similar to *CYP2D7* (172/172 matches, 100% identity) than to *CYP2D6* (164/172 matches, 95% identity). The bar plots again highlight significant differences in introns 2 and 6, which, in this case, align more closely with *CYP2D7*. The alignment results for *CYP2D6* identified three regions that failed to align with the target sequence, two of which were located exclusively in *CYP2D6* introns. These observations confirm that *CYP2D6*68* is a hybrid structure that starts with *CYP2D6* and transitions to *CYP2D7*. Furthermore, in intron 1, the alignment revealed 12 mismatches in the first 144 bp for *CYP2D7*, while *CYP2D6* showed no mismatches in this region. Both genes shared two identical mismatches in other parts of the alignment. These findings confirm that *CYP2D6*68* is a hybrid structure that begins with *CYP2D6* and transitions to *CYP2D7*, with the conversion initiating at intron 1.

## 3 Discussion

Despite its strengths, this study has several limitations. First, potential inaccuracies may persist in the underlying haplotype assemblies. Current long-read sequencing technologies, while powerful, can still incompletely resolve highly complex genomic regions, leading to ambiguities in defining CNVs and other SVs. For instance, confirming a CNV is challenging if the duplicated gene copies are not captured on a single read. Future advancements in long-read accuracy and sequencing depth will be crucial for overcoming these challenges.

Second, our re-annotation of the MHC and KIR loci revealed potential discrepancies in previous annotations, such as misaligned coding sequences and truncated repetitive elements. We attribute this to the ability of TypeAssembly’s local alignment-based method to identify issues missed by minimizer-based aligners like minimap2. However, a systematic investigation is required to pinpoint the exact sources of these previously undocumented errors.

Furthermore, the SV analysis was constrained by its reliance on PharmVar definitions, which limited our resolution to the exon/intron level. This granularity may be insufficient for certain genes and applications. Additionally, the standard VCF format is not well-suited for representing the complex allelic structures identified by our method. This highlights the need for customized SV information for specific gene families, and we need to store it separately from small indels.

Finally, TypeAssembly is computationally intensive compared to minimizer-based algorithms. Its high accuracy is derived from its comprehensive use of BLAST to align all potential genomic and coding sequence alleles to the haplotype assemblies. This exhaustive approach, particularly in mode-FASTA, creates a significant computational bottleneck. The current implementation in Python has also not been optimized for performance, presenting a clear avenue for future engineering and improvement to enhance the tool’s scalability and usability.

## 4 Conclusions

This study develops a haplotype assembly annotation tool, TypeAssembly, and demonstrates its capability to accurately annotate complex genes, including pharmacogenes, *KIR*, and the MHC gene locus. The dual modes, mode-FASTA and mode-VCF, enable precise allele typing for diverse genetic structures and allele definitions. Benchmarking results validated TypeAssembly’s accuracy for 41 genes in the MHC gene locus and 17 *KIR* genes, aligning closely with consensus annotations from Immuannot. Additionally, the annotations provided for 15 pharmacogenes in haplotype assemblies underscore TypeAssembly’s significant contribution to pharmacogenomics research. Through comparison with short-read sequencing tools such as DRAGEN and Aldy, we highlighted the reliability of TypeAssembly and revealed the limitations of short-read approaches in resolving complex genetic structures, reaffirming the value of high-quality haplotype assemblies. TypeAssembly not only establishes the first annotations for pharmacogenes in haplotype assemblies but also serves as a scalable framework for evaluating the accuracy of genome assemblies and annotating genes with well-defined alleles. These contributions hold significant potential for advancing pharmacogenomics research and improving our understanding of genes with complex architectures.

## 5 Methods

### 5.1 Dataset

We analyzed 94 haplotype assemblies from 47 individuals of the HPRC first release, which were generated using publicly available human pangenome production workflows. Allele definitions for 15 genes were obtained from PharmVar (Table 1), with allele sequences and variant information stored in VCF files. To ensure data consistency, we normalized the PharmVar VCF files based on their reference sequences. Definitions of copy number variations and structural variations were also sourced from PharmVar. Among the 15 genes, five exhibited potential for CNVs, while seven were associated with structural variations. For structural variant calling, we utilized coding region sequence definitions provided by GenBank [15] and made each exon and intron sequence correspondingly.

**Table 1:** The list of PharmVar (v6.2.15) genes with details on allele variations.

| Gene | Copy Number Variation | Structural Variation | Numbers of Alleles (Main Alleles) |
| --- | --- | --- | --- |
| *CYP1A2* |  |  | 123 (47) |
| *CYP2A6* | V | V | 126 (45) |
| *CYP2A13* |  |  | 29 (9) |
| *CYP2B6* | V | V | 156 (46) |
| *CYP2C8* |  |  | 54 (17) |
| *CYP2C9* |  |  | 202 (87) |
| *CYP2C19* |  | V | 118 (35) |
| *CYP2D6* | V | V | 551 (170) |
| *CYP3A4* |  |  | 102 (44) |
| *CYP3A5* |  |  | 35 (5) |
| *CYP4F2* | V | V | 64 (21) |
| *DPYD* |  | V | 459 variants |
| *NAT2* |  |  | 154 (58) |
| *NUDT15* |  |  | 50 (20) |
| *SLCO1B1* | V | V | 118 (46) |

**Table 2:** Evaluation of DRAGEN and Aldy pharmacogenomics callers for annotating short-read data.

| Gene | DRAGEN |  |  |  | Aldy |  |  |  |
| --- | --- | --- | --- | --- | --- | --- | --- | --- |
|  | Match | Mismatch | DB Diff. | Total | Match | Mismatch | DB Diff. | Total |
| *CYP2D6* | 84 | 3 | 1 | 88 | 82 | 6 | 0 | 88 |
| *CYP2B6* | 85 | 3 | 0 | 88 | 81 | 7 | 0 | 88 |
| *CYP2C9* | 88 | 0 | 0 | 88 | 88 | 0 | 0 | 88 |
| *CYP2C19* | 88 | 0 | 0 | 88 | 79 | 9 | 0 | 88 |
| *CYP3A5* | 88 | 0 | 0 | 88 | 88 | 0 | 0 | 88 |
| *CYP4F2* | 65 | 8 | 9 | 82 | 77 | 2 | 9 | 88 |
| *NUDT15* | 88 | 0 | 0 | 88 | 88 | 0 | 0 | 88 |
| *SLCO1B1* | 86 | 2 | 0 | 88 | 84 | 4 | 0 | 88 |

### 5.2 mode-FASTA

#### 5.2.1 Preprocessing

In mode-FASTA, TypeAssembly first builds a BLAST database [16] from each haplotype assembly. It then queries these databases using reference gene sequences to locate potential gene clusters, such as *KIR* and MHC. Any resulting hit with an alignment length greater than 50% of the reference gene is kept, and the corresponding region in the assembly is extracted as a target sequence for further analysis.

Next, TypeAssembly performs a more detailed alignment against these extracted target sequences. It uses all known allele genome sequences and all known allele coding sequences as queries. For this BLAST search, the parameters are adjusted to be more tolerant of structural variants by lowering the gap open penalty to 1 and the gap extension penalty to 2. This encourages BLAST to report a single, longer alignment spanning an indel, rather than fragmenting the alignment into two separate parts around the gap.

#### 5.2.2 Allele Typing and Novel Allele Detection

Allele typing employs a hierarchical strategy to classify alleles while allowing for copy number variation. Each tier identifies a specific type of allele, and if an allele is not found in the reference database, it is classified as novel and appended with a special character to denote its type. The first tier identifies perfect matches: full-length genome hits that have 100% identity to an allele in the reference database. These represent standard, unambiguous calls and are reported with their official nomenclature (e.g., *KIR2DL1* *0010101).

Next, a novel non-CDS variant allele ($) is identified when an allele contains variations only in non-coding regions (e.g., introns or UTRs) but its coding sequence (CDS) is identical to a known allele. The process first uses an imperfect full-genomic alignment to anchor the gene region, then stitches together individual exon alignments that match a known allele’s CDS with 100% identity. To be successful, the exon alignments must be consecutive and cover the entire length of the reference CDS. These novel alleles are reported using the known synonymous coding sequence level allele name followed by a $ suffix (e.g., *KIR2DL1* *00101$). A CDS-only allele (+) is identified with the same stitching technique, confirming that disjoint CDS alignments can be assembled to perfectly cover a known reference CDS that is otherwise absent from the full-length genomic database and also reported using the known synonymous coding sequence level allele name (e.g., *KIR2DL4* *00504+).

The hierarchy then proceeds to detect novel alleles containing one or more synonymous variants within the coding sequence (=). The method extends the stitching logic to include CDS alignments with mismatches. For each mismatch, an in-silico translation of the query and subject DNA subsequences is performed. If the resulting amino acid sequences are identical, the variant is confirmed as synonymous. Also, since the query CDS sequence would be strictly divided by three, we only need to move to the nearest codon start position without considering the frameshift. These novel alleles are reported using the allele name truncated to the protein-level identifier, followed by a = suffix (e.g., a novel synonymous variant of the *KIR2DL1* *001 protein would be reported as *KIR2DL1* *001=).

If an allele cannot be classified by any of the prior tiers, a final best-effort assignment (#) is made. This tier considers all remaining full-length genomic alignments with *>*97% identity to a known allele and chooses the one with the lowest normalized edit distance (calculated as edit distance/alignment length) as the most similar match. This classification provides the closest available reference and is reported with a # suffix (e.g., *KIR2DL1* *0010101#). In summary, this mode-FASTA pipeline performs comprehensive copy number estimation, allele typing, and novel allele detection for structurally complex gene families that use FASTA files to define the alleles like *KIR* and MHC.

### 5.3 mode-VCF

#### 5.3.1 Copy Number Estimation

For mode-VCF, TypeAssembly begins by constructing a BLAST database for each haplotype assembly. Using the reference gene sequences as queries, TypeAssembly identifies potential target gene regions in haplotype assemblies. While local alignment is highly efficient, it does not guarantee complete end-to-end coverage of the target sequences. To verify that the BLAST hits correspond to the prospective regions, TypeAssembly applies filters to require a sequence identity greater than 98% and an alignment length exceeding one-third of the reference sequence length. BLAST hits that satisfy these criteria are classified as prospective regions. These regions are subsequently used to assemble full-length target sequences, enabling the determination of copy numbers. Instances of full gene deletion are identified when no BLAST hits meet the defined criteria. The resulting target sequences are then extracted for further analysis.

#### 5.3.2 Small Variant Calling

TypeAssembly aligns target sequences to the reference gene sequence. It initially uses BLAST hits to identify variants if the target sequences are covered end-to-end. However, when the target sequences originate from two separate BLAST hits, typically due to poor alignment in the middle regions or unsuccessful alignment at the ends, TypeAssembly employs an end-to-end dynamic programming-based global alignment on uncovered regions. This alignment utilizes the same scoring function as BLAST: match +1, mismatch *−* 2, gap existence 0, gap extension *−* 2.5. Once TypeAssembly aligns the full sequences to the reference sequences, it identifies single nucleotide polymorphisms (SNPs) where mismatches occur. For cases involving insertions or deletions (indels), TypeAssembly combines SNPs to define the indels. Finally, it normalizes the variant information using bcftools based on the reference sequence and stores the data in VCF files for subsequent analysis.

#### 5.3.3 Structural Variant Calling

Using the exon and intron sequences from both genes and pseudogenes, TypeAssembly aligns each sequence against the target sequences. For gene conversions, this alignment allows TypeAssembly to identify regions of each gene that are more similar to either the original gene or the pseudogene, as well as the starting points of the conversions. In cases of partial gene deletions, it identifies the specific exon or intron regions that fail to align. TypeAssembly then lists the possible alleles where the structural variant information matches the definitions provided by PharmVar.

#### 5.3.4 Allele Typing

TypeAssembly integrates information obtained from small variant calling and structural variant calling to compile a list of possible alleles where the variant or structural variant data matches the definitions provided by PharmVar. To refine this list, TypeAssembly removes definitions that are subsets of others. For instance, the definition of *CYP2D6*10* includes rs1065852 and rs1135840 variants, whereas *CYP2D6*39* includes only rs1135840 variants. In cases where both variants are identified, TypeAssembly considers *CYP2D6*39* redundant as it is encompassed within *CYP2D6*10*. This process enables TypeAssembly to finalize allele determination. It provides users with a complete list of possible allele types for subsequent clinical annotation and interpretation.

## 6 Abbreviations

CPC: Chinese Pangenome Consortium
CPIC: Clinical Pharmacogenetics Implementation Consortium
HLA: Human Leukocyte Antigen
HPRC: Human Pangenome Reference Consortium
indels: insertions or deletions
KIR: Killer cell Immunoglobulin-like Receptor
MHC: Major histocompatibility complex
SNP: Single-Nucleotide Polymorphism
VCF: Variant Calling Format
WGS: Whole Genome Sequencing

## Declarations

### Ethics approval and consent to participate

Not applicable

### Consent for publication

Not applicable

### Availability of data and materials

The TypeAssembly codebase, along with annotations for 15 pharmacogenes, is available on GitHub.

https://github.com/hongsheng-lai/TypeAssembly.

### Authors’ contribution

H-S.L. conducted the experiments, analyzed the results, and wrote the manuscript. Y-S.L. contributed to the annotation of pharmacogenes. H-F.L. assisted with the annotation of CYP2D6. T-Y.C. supported efforts to try SKIRT on CYP2D6. Y-H.T. contributed to the annotation of HLA. T-K.H. contributed to the annotation of KIR. S-K.L. contributed to the annotation of HLA. J.S.H. and P-L.C. advised on the experimental design and reviewed the manuscript. C-Y.C. provided guidance on the experimental design, as well as writing and reviewing the manuscript.

## Acknowledgements

Not applicable

